# *TEMPRANILLO* homologs in apple regulate flowering time in the woodland strawberry *Fragaria vesca*

**DOI:** 10.1101/2022.02.08.479567

**Authors:** Ata Dejahang, Naeimeh Maghsoudi, Amir Mousavi, Nader Farsad-Akhtar, Luis Matias Hernandez, Soraya Pelaz, Kevin Folta, Nasser Mahna

**Author notes:** Corresponding authors: Nasser Mahna.

## Abstract

Many economically valuable fruit trees belong to the *Rosaceae* family. The long juvenile period of fruit trees makes their breeding costly and time-consuming. Therefore, flowering time regulation and shortening the juvenile phase have been a breeding priority for genetic improvement of fruit tree crops. *TEMPRANILLO* genes act as floral inhibitors in Arabidopsis and so, two orthologs of this gene were isolated here from apple (*Malus domestica*) and functionally analysed in Arabidopsis and woodland strawberry by being overexpressed in both model plants, and partially silenced in strawberry. The overexpression of *MdTEM* genes down-regulated *FvFT1, FvGA3OX1*, and *FvGA3OX2* genes in strawberry. The overexpression lines exhibited delayed flowering, both in terms of days to flowering and the number of leaves at flowering. *RNAi-mediated* silencing of *TEM* resulted in early flowering plants. These plants flowered approximately five days earlier than wild type plants with a lower number of leaves. However, the overexpression of *MdTEM* genes could not functionally rescue the *Arabidopsis* mutants. According to the results, it can be concluded that *MdTEM1* and *MdTEM2* are orthologs of *FvTEM* gene, which play an important role in regulating the juvenile phase and flowering time through controlling *FT* and GA biosynthetic gene expression.

## 1. Introduction

Many economically valuable fruit and ornamental crops belong to the *Rosaceae* family, and while there has been significant progress in the accumulation of genomics-level data in the last decade, the formidable task of linking genes to horticulturally relevant traits remains undone (Folta and Gardiner 2009). The plant’s life cycle is divided into juvenile and adult phases. Plants in the juvenile phase are incapable of response to floral signals even under the inductive conditions (Huijser and Schmid 2011; Yamagishi et al. 2014). Transition from juvenile to adult phase takes a long time in most of the fruit trees (e.g. 5-12 years for apple), which limits and slows breeding efforts such as backcrosses, inbreeding, or production of new hybrids (Flachowsky et al. 2009). Therefore, flowering time regulation is of key importance to breeding programs (Jung 2017). In the past decades, great progress has been obtained in understanding the molecular basis involved in flowering time regulation, especially in *Arabidopsis* (Liu et al. 2017). Several genetic pathways which control flowering, including the vernalization, photoperiod, gibberellins content, autonomous, and the ambient temperature pathways, have been characterized. These signaling pathways integrate developmental and environmental factors associated with the activation of a key floral regulator, *FLOWERING LOCUS T* (*FT*) (Fornara et al. 2010; Wellmer and Riechmann 2010). FT protein is produced in leaves and moves through the phloem to the apex where it forms a complex with a bZIP transcription factor FD and activates expression of the floral meristem identity genes to promote flowering (Corbesier et al. 2007). The fact that plants are unable to initiate flowering during juvenility phase even in inductive environmental conditions proposes that inhibitory mechanisms may suppress FT expression during juvenility and prevent flowering (Corbesier et al. 2007; Sgamma et al. 2014). Many plants require a given day length sometimes in combination with a certain temperature to initiate flowers. Flowering at right time associated with the seasonal and endogenous signals is vital for successful reproduction in plants (Liu et al. 2017). Numerous genes influencing floral induction have been characterized. Orthologs of flowering-related genes of *Arabidopsis thaliana* (L.) have been isolated from rosaceous crops like apple (*Malus* x *domestica*), including *LEAFY (LFY), APETELA1 (AP1), AGAMOUS (AG), TERMINAL FLOWER (TFL1), BpMADS4*, and *SERRATED LEAVES AND EARLY FLOWERING (SEF)* (Kotoda et al. 2002; Wada et al. 2002; Esumi et al. 2005; Kotoda and Wada 2005; Flachowsky et al. 2007; Maghsoudi et al. 2015). *TEMPRANILLO* genes (*TEM1* and *TEM2*) belong to the plant-specific transcription factor RAV (Related to ABI3/VP1) subfamily, which contain two DNA-binding domains, an AP2/ERF and a B3 DNA-binding domain (Castillejo and Pelaz 2008; Matías-Hernández et al. 2014). *TEM* genes play a pivotal role in Arabidopsis flowering time. They directly repress *FT* transcription through binding to two regions in the *FT* gene 5’ untranslated region (Castillejo and Pelaz 2008) and also repress the GA biosynthetic genes *GA3OX1and GA3OX2* through binding to a sequence in the first exon (Osnato et al. 2012; Matías-Hernández et al. 2014). Double mutant *Arabidopsis* plants with reduced *TEM1* and *TEM2* activity flower earlier than the single *tem1* and *tem2* mutants, which flower earlier than wild-type plants (Castillejo and Pelaz 2008).

Plant trichomes are specific epidermal protrusions which have several characteristics that vary between plant species and organs. The juvenile-to-adult transition is correlated with various morphological changes, including the formation of trichomes on the abaxial side of leaves (Wang et al. 2019). Osnato et al. (2012) reported that TEM genes control floral transition through linking the photoperiod and GA-dependent flowering pathways to the regulation of the floral integrators. The timing of abaxial trichome formation is associated with flowering time, consistent with the fact that the juvenile-to-adult vegetative phase change contributes to acquisition of the competence to flower (Aguilar-Jaramillo et al. 2019). Studies revealed that TEM genes inhibit trichome initiation from the mesophyll, the lower layer of epidermis (Jiao 2016; Matías-Hernández et al. 2016). Fluorescently labeled GA3 exclusively accumulated in the mesophyll of cells, but not in the epidermis, suggesting that TEM plays an essential role in GA biosynthesis and distribution in the mesophyll, resulting in the epidermal trichome formation (Matías-Hernández et al. 2016).

*F. vesca* offers several features that makes it an appropriate plant for functional genomics research in the Rosaceae family. It has a small diploid genome that is fully sequenced. The plant are small, easily transformed, maintain small size, self-compatible, ever-bearing, and may be propagated by runners and branch crowns as well as by seed (Folta and Davis 2006; Slovin et al. 2009; Pantazis et al. 2013). *F. vesca* has both seasonal (SD) and perpetual flowering accessions with different photoperiodic responses (Brown and Wareing 1965; Heide and Sønsteby 2007). In perpetual flowering accessions, LD advances flower induction, but plants eventually flower also under SD conditions (Koskela et al. 2012; Rantanen et al. 2014; Koskela et al. 2017). Recently, the full length cDNA of *FvTEM* was isolated and characterized *in-silico*, which revealed that it was 1152 bp in length predicted to encode 383 amino acids and homologous to *AtTEM* (Dejahang et al. 2018). In the present study, *MdTEM1* and *MdTEM2* genes overexpressing and RNAi silencing transgenic strawberry plants were generated for evaluating and studying their role on the regulation of flowering time. The transgenic plants were evaluated for gene integration and expression during *in vitro* cultivation. Flower initiation and development were studied on *in vitro* shoots and/or glasshouse-grown plants.

## 2. Materials and Methods

### 2.1 Plant Material and growth conditions

#### Apple

apple trees cv. Golden Delicious were used for DNA and RNA extractions, gene isolation and gene expression analyses.

#### Strawberry

Seeds from perpetual flowering LD accession Hawaii-4 (H4) of the woodland strawberry, *Fragaria vesca* L. (PI551572), were sterilized for 5 minutes in 70% (v/v) ethanol and in 1% sodium hypochlorite with 2 drops of Tween 20^®^ for 5 min and then rinsed in sterile distilled water several times before germination in Petri dishes containing 1/2MS at pH 5.7 with 3% (w/v) sucrose. Seeds were cultured initially at 22°C in dark for one week, then at 25°C under 16/8 hr. photoperiod. Seedlings with one or more true leaves were transferred into jars containing MS medium to increase size. The seedlings were transferred into fresh medium two weeks before the transformation.

#### Arabidopsis

Seeds of Col-0 (wild type) and *tem* mutants were planted in pots under 16-8 h photoperiod and 21-19 day-night temperature and 60% humidity in a growth chamber.

### 2.2 MdTEM genes isolation and vector construction

Two *AtTEM1*/*AtTEM2* homologs were isolated from leaf samples of 10-year-old trees of Golden Delicious cultivar of apple (*Malus domestica*) (designated as *MdTEM1* and *MdTEM2*) and cloned into *pAlligator2* (Bensmihen et al. 2004) independently under the control of double enhanced *CaMV35* promoter and *NOS* terminator for overexpression. A 137 bp fragment of *MdTEM1* was also cloned into *pHellsgate12* under the control of *CaMV35S* promoter and *octopine synthase* terminator with an intron region designed to trigger RNAi-mediated gene silencing. The *nptII* gene was used as selectable marker under the control of *NOS* promoter and terminator. Vectors carrying overexpression and RNAi constructs were incorporated into *Agrobacterium tumefaciens* strains GV3101 and GV2260 through electroporation (GenePulser, BioRad, USA)

### 2.3 Plant transformation and regeneration

#### Strawberry

For transformation and regeneration of wild strawberry, young fully expanded leaflets were placed adaxial side up in a Petri dish and sliced across and/or along the secondary veins to produce multiple cuts. Leaf sections were co-cultivated with *Agrobacterium* harboring an overexpression or RNAi construct in media containing MS salts and vitamins, 2% sucrose, 3 mg/L BA, 0.2 mg/L IBA and 0.7% agar (Oosumi et al. 2006). After 3 days of co-cultivation, explants were washed with liquid MS containing 500 mg/L cefotaxime and placed abaxial side up in the selection media containing 3 mg/L BA, 0.2 mg/L IBA, 25 mg/L kanamycin and 250 mg/ml cefotaxime. Explants were subcultured two-week interval for 60-90 days until shoots appeared. Transformation efficiency for each construct was calculated as the percentage of number of explants which produced PCR-positive plants out of the total number of inoculated explants.

#### Arabidopsis

*Agrobacterium* harboring *MdTEM1* and *MdTEM2* gene were used to transform different *Arabidopsis tem1tem2, tem1* and *tem2* mutants, and Col-0 as control using the floral dip method (Clough and Bent 1998). Transgenic seeds were selected by GFP expression in their seed coat under a fluorescent microscope.

### 2.4 PCR analysis of transgenic strawberry plants

DNA was isolated from leaves of transformed and untransformed plants using a modified CTAB method (Doyle and Doyle 1990). Quality and quantity of the extracted DNA were checked by agarose gel and NanoDrop1000 spectrophotometer (NanoDrop Technologies, Wilmington, DE, USA). The putative transgenic plants were screened for the presence of T-DNA by polymerase chain reaction (PCR) analysis using NOS terminator and *MdTEM1* and *MdTEM2* primers for overexpression and using *MdTEM1* and *NPTII* primers for RNAi silencing experiments. Primers used for plant transformation validation by PCR are listed in Supplementary Table 1. The PCR reaction was carried out using 100 ng of genomic DNA using the following step cycle program: 94°C, 30 s; 57°C, 30 s, and 72°C, 50s for 32 cycles. A 5μL aliquot of each PCR reaction was analyzed by 1% agarose gel electrophoresis.

### 2.5 RNA extraction, cDNA synthesis, and real-time PCR

Total RNA was extracted from the youngest fully opened leaves using modified CTAB method (Monte and Somerville 2002) and treated with RNase-Free DNase (Fermentas, Germany) according to the manufacturer’s recommendations. The purity and concentration of total RNA were measured using a NanoDrop 1000 spectrophotometer (NanoDrop Technologies, Wilmington, DE, USA) and first strand cDNA synthesized from 500 ng total-RNA using MMLV reverse transcriptase and oligo dT. qRT-PCR reactions were performed in a final volume of 20μl on the Corbett Rotor-Gene 6000 (Corbett LifeScience) using Power SYBR green master mix (Life Technologies). The PCR conditions were as follows: 95°C for 5 minutes, followed by 40 cycles of 95°C for 15 seconds and at 60°C for 35 seconds. Melting-curve analysis was conducted to verify the specificity of each primer using a temperature ramp starting from 65°C to reach 95°C with fluorescence measured every 1°C. All qRT-PCRs were run in two technical and three biological replicates.

Relative transcript levels of *MdTEM1, MdTEM2* and *MdFT* genes from apple as well as *FvTEM, FvFT, FvGA3OX1*, and *FvGA3OX2* genes from strawberry were calculated by the 2 ^ΔCt^ for apple genes and 2^−ΔΔCt^ and −1/2^−ΔΔCt^ for up and down-regulated genes in wild strawberry, respectively (Schmittgen and Livak 2008). *MdActin* for apples genes and *FvMSI1* for wild strawberry genes were used as constitutive reference genes. Primers used for qRT-PCR analysis are listed in Supplementary Table 1.

### 2.6 Flowering time analysis

For strawberry plants, independent transgenic lines and WT plants were rooted, transferred into pots and grown in the phytotron. Soilless growing media consisted of fertilized peat supplemented with 25% (v/v) of vermiculite. Flowering was recorded as the date of the first open flower. Flowering time differences between H4 and transgenic lines were observed by counting the number of days before flowering and the number of leaves in the primary leaf rosette before flowering.

For *Arabidopsis* plants, the transgenic seeds were grown up to T2 generation to have enough seeds for evaluating flowering time. Rosette and cauline leaves were counted as an index for flowering time.

### 2.7 Trichome analysis

In strawberry, four independent overexpressed lines (35S::*MdTEM1*#1 and 35S::*MdTEM1*#2, 35S::*MdTEM2*#1 and 35S::*MdTEM2*#2), and three independent silenced lines (RNAi-TEM #1, #2 and #3) as well as the WT plants with three biological replicates were studied. The fully expanded leaves from each line were placed in glass flasks containing 50 ml of 70 % ethanol. Paradermal sections were collected from the central region of the abaxial surface of these adult leaves. The sections were washed using sterile distilled water for 3 min and immersed in 10 % sodium hypochlorite solution until total clearing. The sections were then washed in distilled water and stained with 1% safranin for 3 h and rinsed in distilled water to remove excess dye. For each line, a total of 3 slides containing 5 sections each were prepared, making a total of 15 sections per line. The non-glandular trichomes were scored using a stereomicroscope equipped with a 14x objective lens (Figueiredo et al. 2013).

### 2.8 Phylogenetic analyses

Multiple sequence alignment was performed using the deduced amino acid sequences of *MdTEM1, MdTEM2* and *FvTEM* with other *RAVI* orthologs from different plants. The sequences were aligned using the CLUSTALW alignment tool in MEGA7. The phylogenetic tree was constructed with the maximum parsimony method and visualized using MEGA7. Bootstrap values were estimated using 500 replications. Numbers at each node indicate the percentage of bootstrap samples.

### 2.9 Statistical analysis

ANOVA was conducted on the averages using the general linear model, and differences between means were analyzed by LSD test. All statistical analyses were conducted using the SPSS software package version 16.0 (SPSS, Inc., Chicago, Illinois).

## 3. Results

### 3.1 Domain structure and phylogenetic analysis

The full-length cDNA of *MdTEM1, MdTEM2* and *FvTEM* consisted of the coding sequences of 1221, 1206 and 1152 bp, respectively, were isolated, predicted to encode a protein with 406, 401 and 383 amino acids, respectively. They had no intron region and consisted of the AP2 and B3 domains which characterize it as a member of the RAV1 protein family. *In-silico* comparison of *FvTEM, MdTEM1* and *MdTEM2, AtTEM1* and *AtTEM2* showed that they shared a high homology, and all have AP2 and B3 conserved domains (Supplementary Fig. 1a). To determine the evolutionary relationships among the RAV1 family proteins, phylogenetic analysis was conducted by the amino acid sequences using Neighbor–Joining method for generating the phylogenetic tree. Phylogenetic analysis demonstrated that *MdTEM1, MdTEM2* and *FvTEM* are homologous to RAV1 family proteins from other plants (Supplementary Fig. 1b).

### 3.2 Relative expression of MdTEM genes in apple

The relative steady-state transcript levels of *MdTEM1, MdTEM2* and *MdFT* genes were measured in different tissues of apple by RT-pPCR. The highest expression levels of *MdTEM1* were observed in juvenile leaves and roots (Fig. 1), while the lowest transcript accumulation levels were obtained in flowers and mature stems. However, *MdTEM2* showed an almost opposite expression pattern, the highest expression levels were found in mature stems, flowers and fruits, whereas the lowest were observed in juvenile leaves and roots. On the other hand, *MdFT* had higher relatively expression in fruits, flowers and mature stems.

**Fig. 1.**
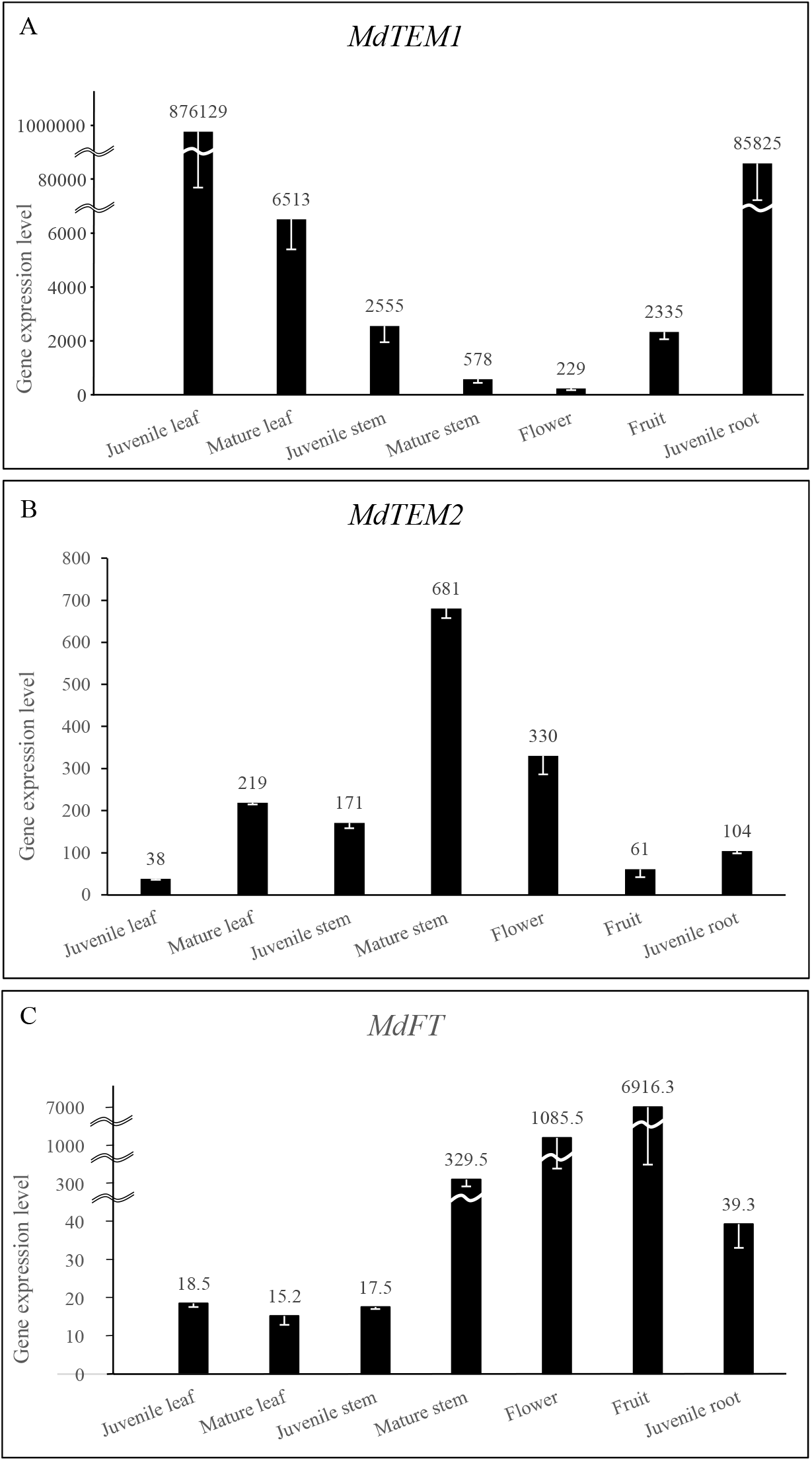
Expression levels of A) *MdTEM1*, B) *MdTEM2*, and 3) *MdFT* genes in different tissues in apple measured by qRT-PCR. Error bars are showing standard errors.

### 3.3 Regeneration and transformation validation

Two over-expression and one RNAi silencing constructs were introduced into the diploid strawberry using *A. tumefaciens* GV3101. Putative transformants regenerated and rooted on MS medium containing kanamycin (Supplementary Fig. 2). The resistant plantlets and non-transformed controls were then grown in pots in a phytotron. Regenerated plants were screened by PCR using *MdTEM1* and *MdTEM2* and NOS terminator primers for both over-expression constructs and by *MdTEM1* and *NPTII* primers for *RNAi* silencing construct. PCR analysis revealed the amplification of the expected specific fragments in transformed plants. No amplification was detected in the non-transgenic control (Supplementary Fig. 3). Transformation efficiency was calculated based on the percentage of inoculated explants that resulted in the production of PCR-positive plants. The results showed that the efficiency of transformation for *35S::MdTEM1, 35S::MdTEM2* and RNAi-*MdTEM1* constructs were 23.57, 17.07 and 32.5% respectively (Supplementary Table 2).

### 3.4 Gene expression analysis

To study the role of *TEMPRANILLO* as a flowering-related transcription factor, the expression of *FvFT1, FvTEM, FvGA3ox1* and *FvGA3ox2* were measured by Real-Time quantitative PCR in the transgenic lines and non-transgenic control plants. qPCR analysis showed that altered expression of *TEM* could affect the transcript levels of floral integration genes. It was confirmed that RNAi-*MdTEM1* lines exhibited lower *FvTEM* transcript levels. The RNAi-*MdTEM1* lines showed significant increased transcript accumulation of *FvFT1, FvGA3ox1* and *FvGA3ox2* compared to control plants. Overexpression of *MdTEM* lines exhibited a significant decrease in *FvFT1, FvGA3ox1* and *FvGA3ox2* transcript accumulation compared to WT plants (Fig. 2). However, the results also revealed that the *MdTEM1* and *MdTEM2* had no effect on endogenous *FvTEM* expression, although they were successfully expressed in 35S::*MdTEM1* and 35S::*MdTEM2* lines with different expression level (Fig 3).

**Fig. 2.**
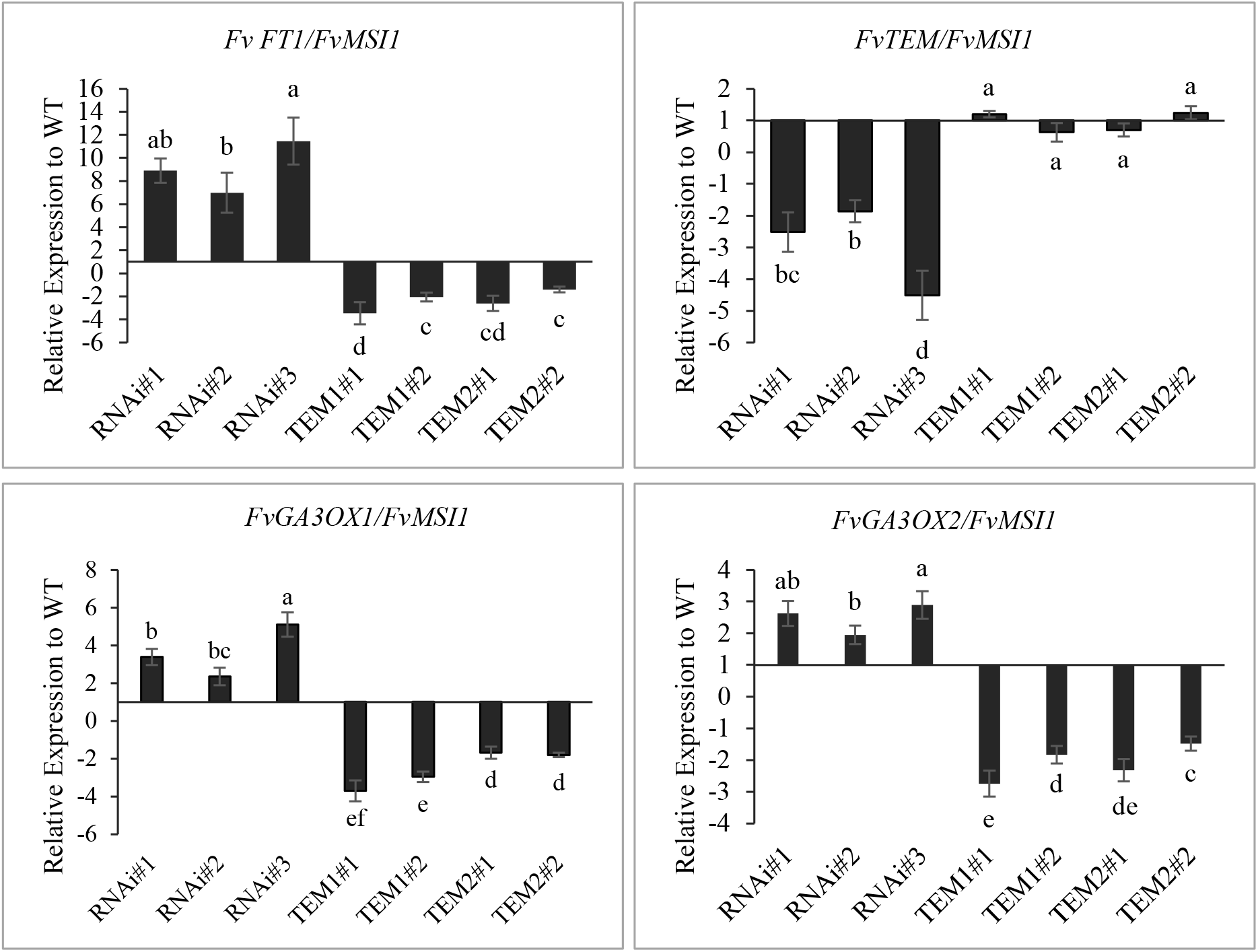
Relative expression of *FvFT1, FvTEM, FvGA3ox1* and *FvGA3ox2* genes in overexpressing and silencing lines. The data are shown as the mean ±SE of three biological and three technical replicates. Data are presented relative to the H4 control plant for each transgenic construct (three silencing lines and two overexpressed lines for each TEM1 and TEM2 constructs.

**Fig. 3.**
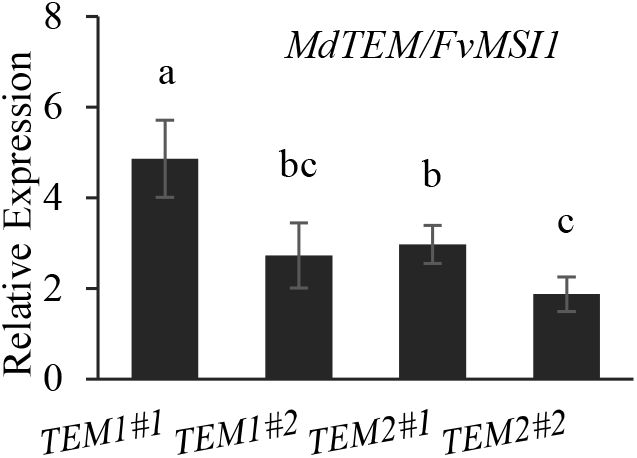
Relative expression of *MdTEM1* and *MdTEM2* genes in over-expressing *35S::MdTEM1* and *35S::MdTEM2* lines

### 3.5 Flowering time analysis

Flowering time was analyzed in the silenced and over-expressed lines compared with non-transgenic controls. The number of leaves before flowering and the number of days to flowering was assessed. The results indicated that the overexpression of *MdTEM1* and *MdTEM2* delayed flowering in *Fragaria vesca*, while the *FvTEM* produced by RNAi-*MdTEM1* silencing lines significantly accelerated flowering (P< 0.01; Supplementary Table 3). Conversely, 25% of RNAi-*MdTEM1* plants were flowered after 30 days when *MdTEM* over-expressing and control plants remained vegetative (Fig. 4a). After 40 days, the percentage of 35S::*MdTEM1* and 35S::*MdTEM2* flowering plants were 36.4% and 37.5%, respectively, whereas this value for silencing RNAi-*MdTEM1*and control plants were 100% (Table 1).

**Table 1.**
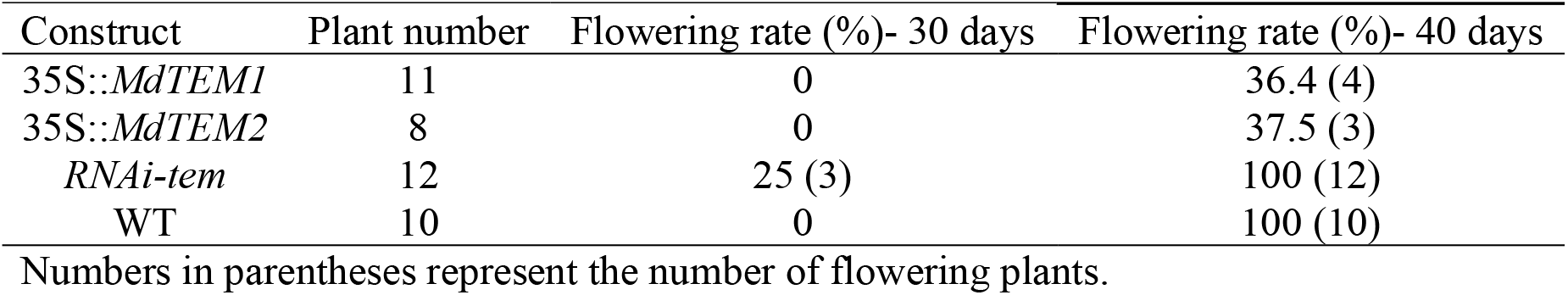
Comparison of flowering rate in over-expressed and silenced lines after 30 and 40 days

**Fig.4.**
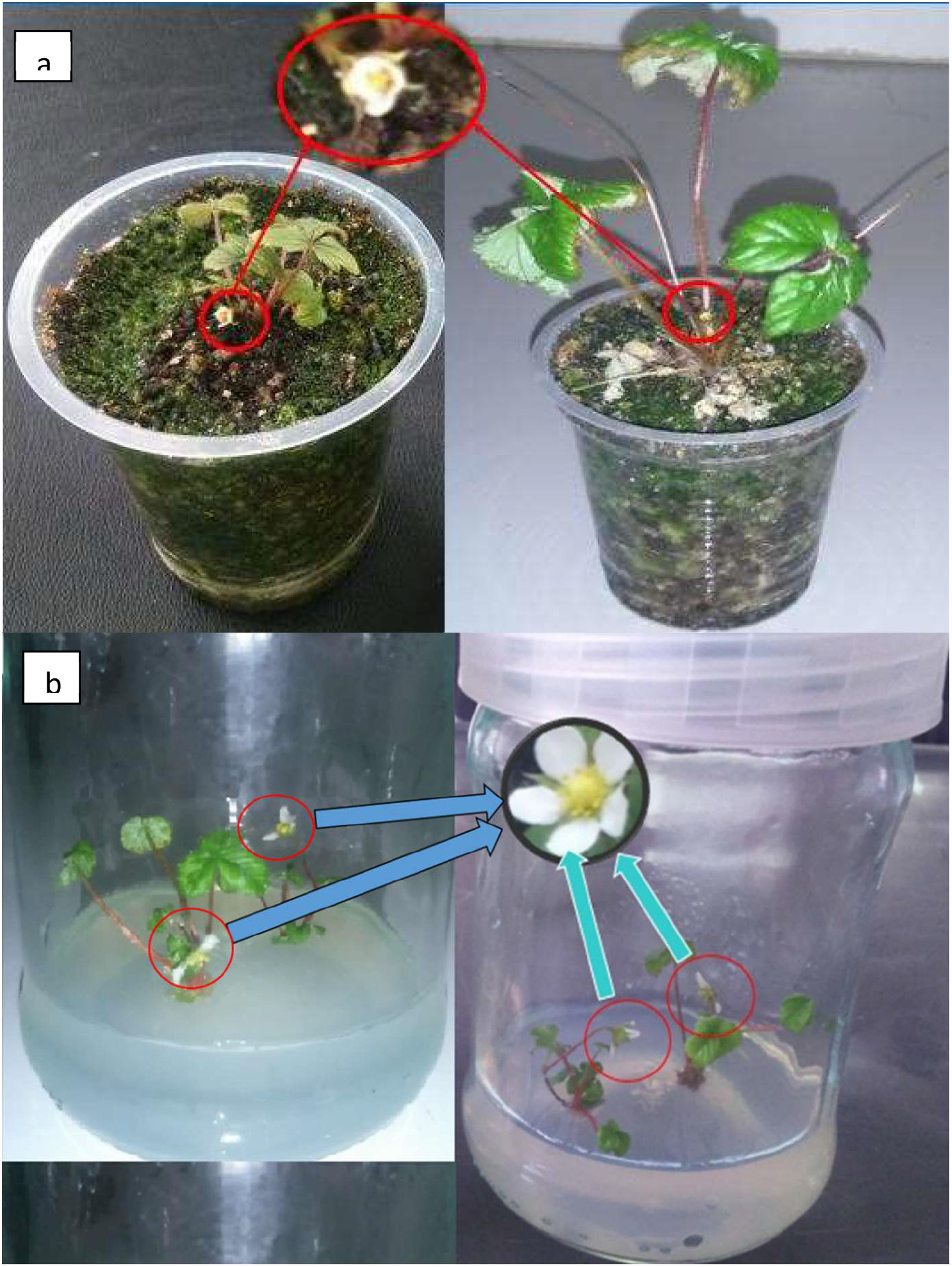
Early flowering in *RNAi*-tem silencing lines a) under inductive conditions in growth chamber and b) under *in-vitro* un-inductive conditions.

In strawberry, overexpression of *MdTEM* caused delayed flowering, denoted as an increase in the number of leaves and the number of days upon flowering. The average number of leaves before flowering for RNAi-*MdTEM1*and control plants were 4.25 and 6.88, respectively. While this value for the *35S::MdTEM1* and *35S::MdTEM2* lines were 11.25 and 12.67, respectively (Fig. 5). The average number of days before flowering for *35S::MdTEM1* and *35S::MdTEM2* were 46.5 and 43.67, respectively, while RNAi-*MdTEM1* lines and control plants flowered after 32.17 and 36.88 days in inductive conditions, respectively. Our results showed that the RNAi-*MdTEM1* silencing plants flowered with a significant smaller number of leaves and days before flowering compared to control plants. Strikingly one of the RNAi-*MdTEM1* lines flowered under in-vitro non-inductive conditions (Fig. 4b).

**Fig. 5.**
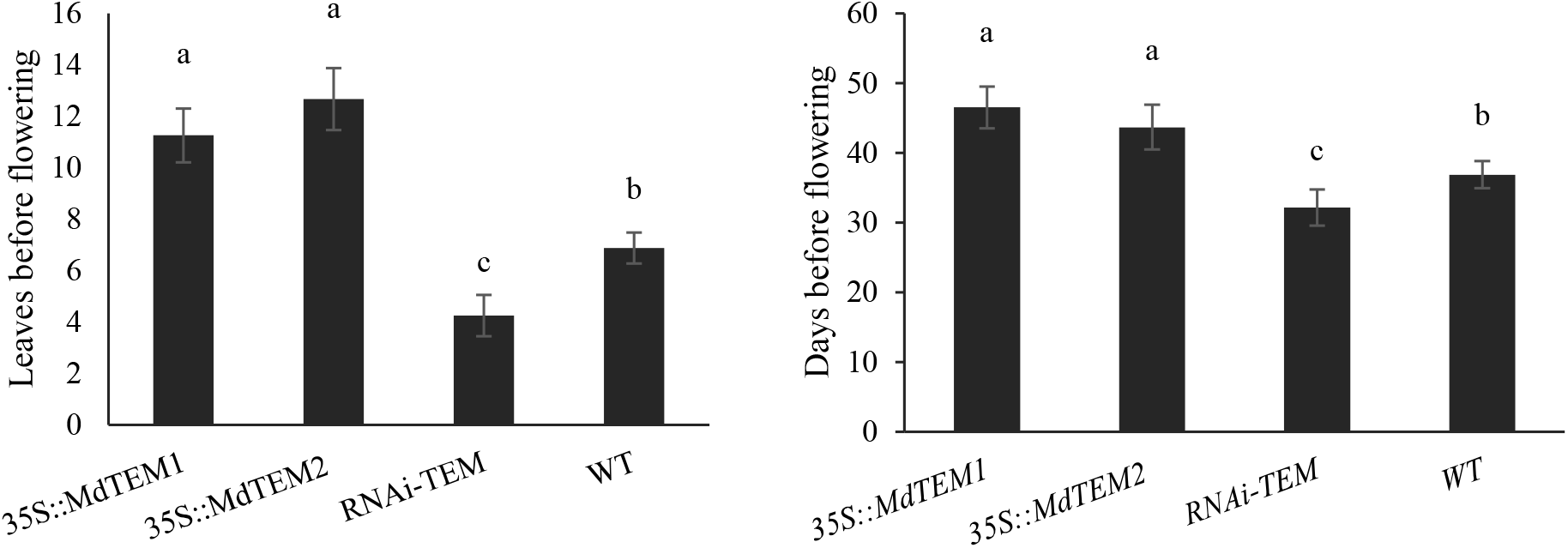
The number of leaves and days before flowering in *MdTEM* overexpressed, silenced and wild type control plants of *Fragaria vesca*.

The effects of *MdTEM1* and *MdTEM2* overexpression on the flowering time was investigated in wild type and different *Arabidopsis* mutants. The results showed that the wild type *Arabidopsis* overexpressed with *MdTEM1* and *MdTEM2* (WT* MdTEM1-T and WT* MdTEM2-T) genes flowered after 14.3 and 14.0 rosette leaves, respectively, which was not significantly different than the wild-type comparator with 14.5 rosette leaves (Table 2). Moreover, the rosette leaf number for *tem1/2* double mutant overexpressed with *MdTEM* genes were same as the *tem1/2* mutant, and the same results were obtained for *tem1* and *tem2* mutants. Overall, our results indicated that the *MdTEM* genes could not functionally complement the *AtTEM* roles in the *Arabidopsis* mutants.

**Table 2.**
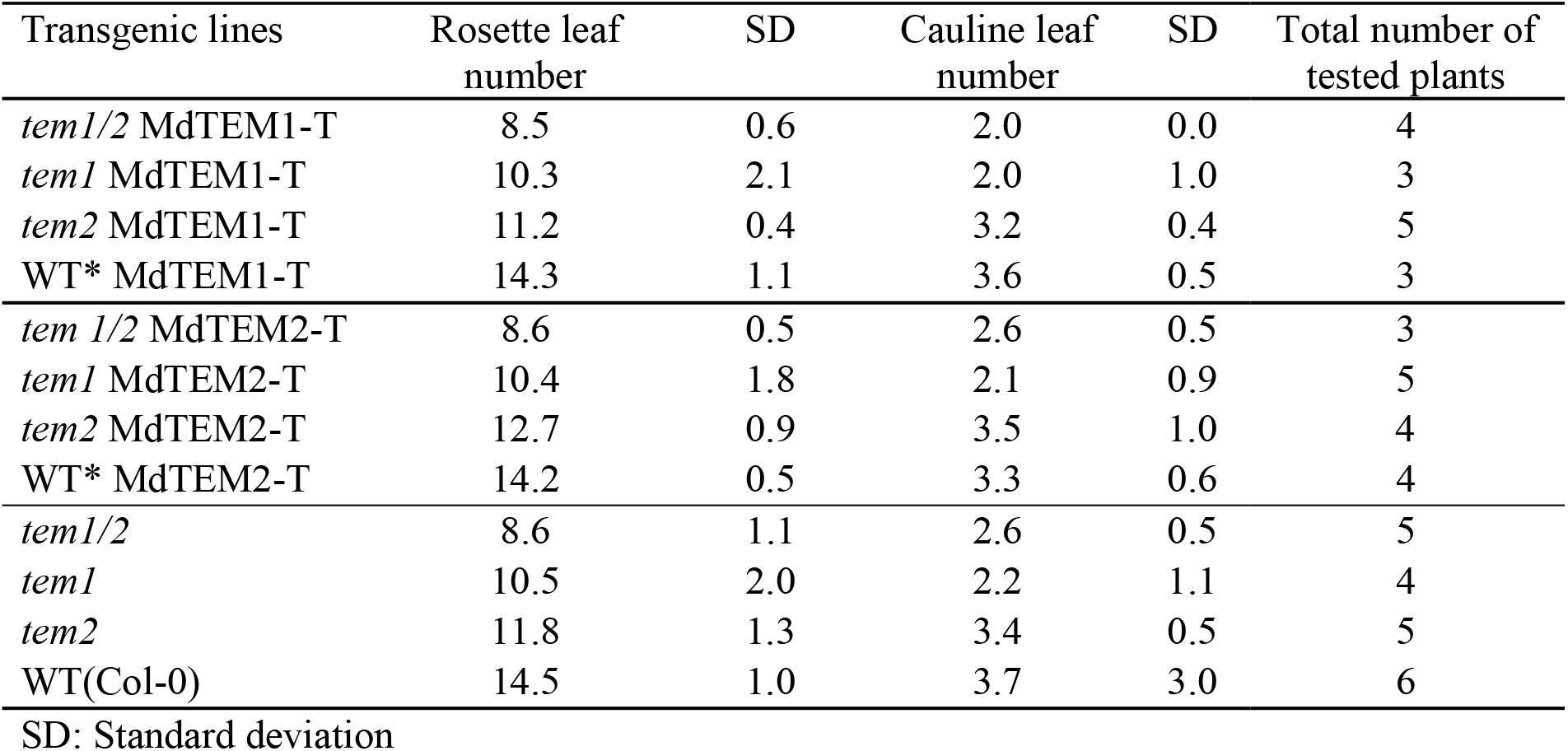
Flowering time assessment in *Arabidopsis* plants

| Transgenic lines | Rosette leaf number | SD | Cauline leaf number | SD | Total number of tested plants |
| --- | --- | --- | --- | --- | --- |
| *tem1/2* MdTEM1-T | 8.5 | 0.6 | 2.0 | 0.0 | 4 |
| *tem1* MdTEM1-T | 10.3 | 2.1 | 2.0 | 1.0 | 3 |
| *tem2* MdTEM1-T | 11.2 | 0.4 | 3.2 | 0.4 | 5 |
| WT* MdTEM1-T | 14.3 | 1.1 | 3.6 | 0.5 | 3 |
| *tem1/2* MdTEM2-T | 8.6 | 0.5 | 2.6 | 0.5 | 3 |
| *tem1* MdTEM2-T | 10.4 | 1.8 | 2.1 | 0.9 | 5 |
| *tem2* MdTEM2-T | 12.7 | 0.9 | 3.5 | 1.0 | 4 |
| WT* MdTEM2-T | 14.2 | 0.5 | 3.3 | 0.6 | 4 |
| *tem1/2* | 8.6 | 1.1 | 2.6 | 0.5 | 5 |
| *tem1* | 10.5 | 2.0 | 2.2 | 1.1 | 4 |
| *tem2* | 11.8 | 1.3 | 3.4 | 0.5 | 5 |
| WT(Col-0) | 14.5 | 1.0 | 3.7 | 3.0 | 6 |

SD: Standard deviation

### 3.6 Trichome analysis

To evaluate the effect of *MdTEM* on the trichome formation in diplod strawberry, the trichome distribution on the abaxial side of the four overexpressed, and three silenced lines as well as the wild type strawberry plants were studied. The results showed that the expression level of *MdTEM* had significant effect on the number of trichomes per mm^2^ (T mm^−2^) on the abaxial side of strawberry leaves. Number of trichomes per mm^2^ in RNAi-*MdTEM1* and *35S::MdTEM* lines were higher and lower than the wild-type control plants, respectively (Fig. 6). The highest number of trichomes per mm^2^ (35.6) was belonged to the RNAi-TEM #1 line, while the 35S::*MdTEM1*#1 line produced the lowest trichomes number (3.33 T mm^−2^). These differences were also observed in microscopic analysis of the abaxial side of their leaves (Fig. 7).

**Fig. 6.**
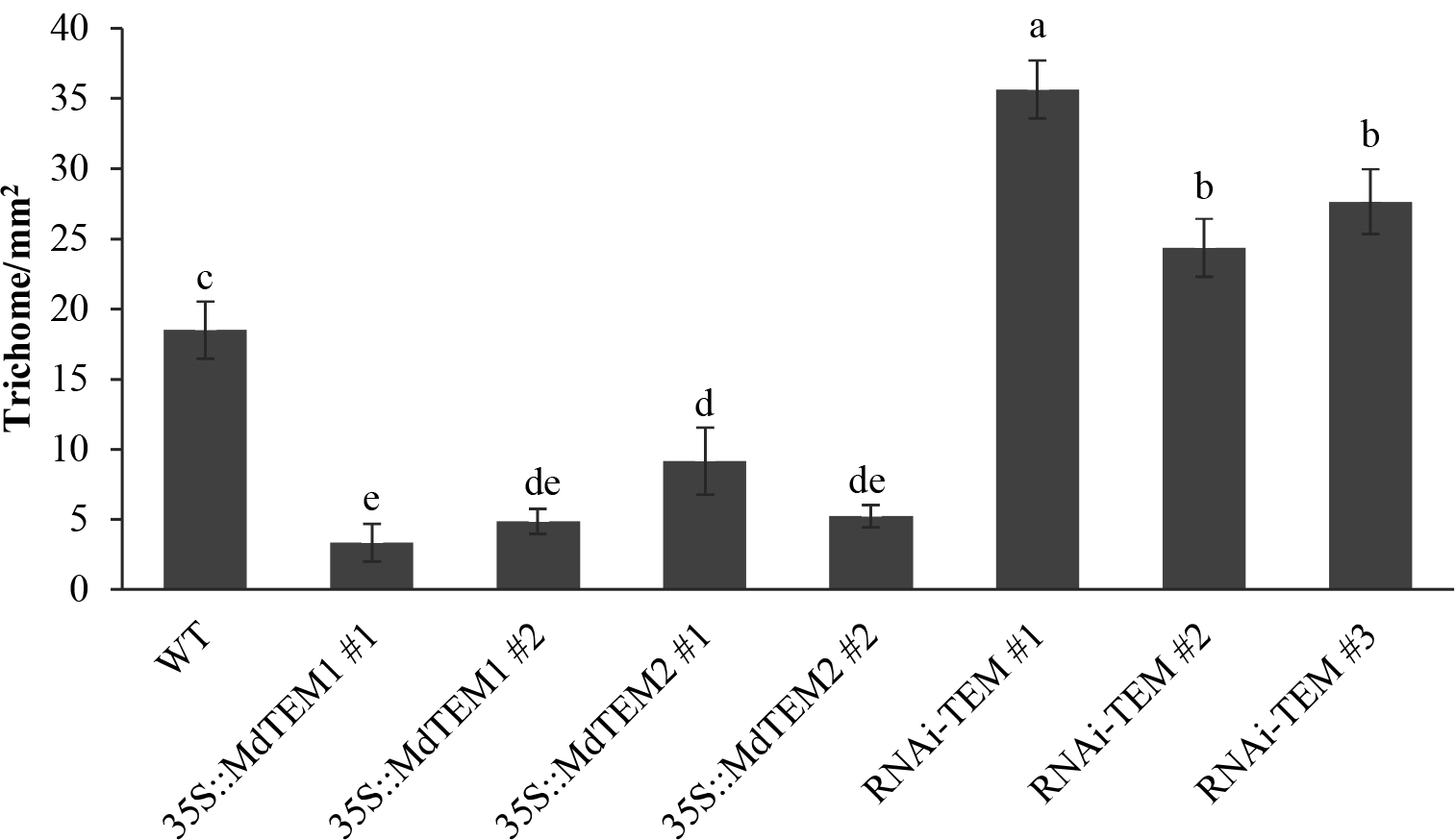
Number of trichomes per mm^2^ in different *MdTEM* overexpressed, silenced and wild type control plants of *Fragaria vesca*. Different letters show significant difference at p≤0.5. Error bars are showing standard errors.

**Fig. 7.**
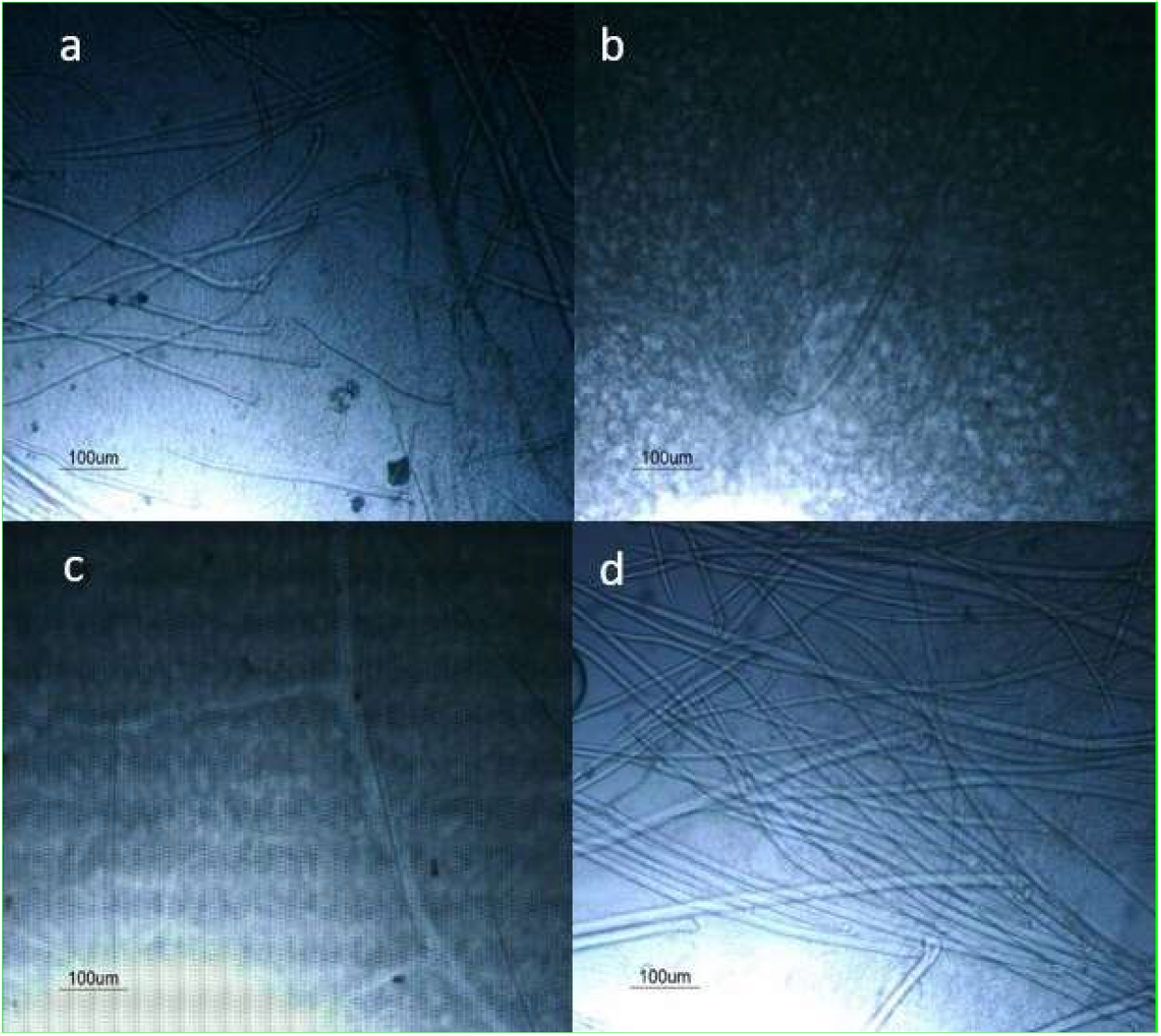
Distribution pattern of trichomes on the abaxial side of *Fragaria vesca* transgenic lines leaves. a) Control b) 35S::*MdTEM1* c) 35S::*MdTEM2* d) RNAi-*TEM*

## 4. Discussion

Development of tree crops by breeding programs is very slow due to their long juvenile phase, which may take several years. One of the most important priorities of most breeding programs is to reduce the juvenile phase and accelerate the flowering process. Decreasing juvenility through use of genetic engineering methods may hasten the production of new cultivars that are desperately needed to meet contemporary challenges, such as changes in climate and pest/pathogen threats. In this study, we produced early flowering lines by down-regulation of *FvTEM-like* genes through RNAi-mediated gene silencing in diploid strawberry, an appropriate model for apple and other species in the *Rosaceae* family. This strategy can be used in other fruit trees with a long juvenile phase. *TEM1* and *TEM2* belong to RAV transcription factors family, have been recognized as flowering repressors and juvenility regulator. Our previous study identified *FvTEM* as a RAV family member, which contained B3 and AP2 domains and can act as floral repressors (Dejahang et al. 2018).

In the present study, we produced early flowering strawberry plants which flowered approximately 5 d before non-transformed control plants. Recently, early flowering strawberry plants produced using ALSV vector containing the *Arabidopsis thaliana FT* gene (Li et al. 2019) and using the *Eriobotrya japonica LEAFY* gene (Liu et al. 2017) have been described. In the present study, the overexpression of *MdTEM1* and *MdTEM2* delayed flowering time in *Fragaria vesca*, indicating that these genes have similar functions to *AtTEM* genes. It has been described that FT protein transported from leaf to the apex and activates downstream genes such as *SOC1, LFY*, and *AP1*, resulting in flower induction (Corbesier et al. 2007). Reciprocal relationships between *TEM* with *FT* and *GA3OX1* and *GA3OX2* were observed, with increasing in mRNA level of *TEM* in overexpressed lines, the level of *FT* and *GA3OX1* and *GA3OX2* were decreased and caused delayed flowering. Also, down-regulation of *TEM* in RNAi-*MdTEM1* lines resulted in an increased level of *FT* and *GA3OX1* and *GA3OX2* transcripts, leading to early flowering. These results were in a good agreement with previous studies that characterize *TEM* as a floral repressor (Castillejo and Pelaz 2008; Osnato et al. 2012; Sgamma et al. 2014; Marín-González et al. 2015). Osnato et al. (2012) reported that constitutive overexpression of *TEM1* in *Arabidopsis* resulted in down-regulation of *GA3OX* genes by binding to their first exon, whereas *tem1-1* and *tem1–1 tem2– 2* mutants showed an up-regulation in *GA3OX1* and *GA3OX2* expression. Taken together with the above observations, we concluded that *MdTEM* could complement the floral repressor function of *FvTEM*, and the expression level of *FvTEM* determine the flowering time through *FT* and GA biosynthetic genes expression in strawberry. On the contrary, our results revealed that the WT *Arabidopsis* overexpressed with *MdTEM1* and *MdTEM2* genes flowered with 14.3 and 14.0 rosette leaves, respectively, similarly to the Col-0 control plants that flowered with 14.5 rosette leaves. However, the rosette leaf number for *tem1/2* double mutant overexpressed with *MdTEM* genes were same as the *tem1/2* mutant, and the same results were obtained for *tem1* and *tem2* mutants. Taken all together, it can be concluded that the *MdTEM* genes could not functionally complement the *AtTEM* roles and also couldn’t rescue the *tem Arabidopsis* mutants. While, *MdTEM* genes significantly delayed the flowering process in wild strawberry, which was in a good agreement with the results of Sgamma et al. (2014) which showed that *TEM* orthologue from *Antirrhinum majus* (*AmTEM*) functionally complemented the role of *AtTEM1* in the *tem1 Arabidopsis* mutant and postponed the transition process from the juvenile to adult phase. These contrasting results on the role of *MdTEM* genes in strawberry and *Arabidopsis* could be attributed to the specific transcription regulatory elements in the systematically related species, as both strawberry and apple belong to the *Rosaceae* family.

The results revealed that the highest number of trichomes per mm^2^ was observed on the abaxial side of RNAi-*MdTEM* and this value for control plants was higher than the *35S::MdTEM* lines. These results indicated that *TEM1* and *TEM2* not only suppress the floral induction but also inhibit the trichome formation through the GA biosynthesis pathway (Matías-Hernández et al. 2014; Matías-Hernández et al. 2016). In fact, flowering is not the only process controlled by RAV proteins, but has also been found to be involved in other plant growth processes such as trichome formation, leaf senescence, and responses to pathogenic infections, abiotic stresses (Wang et al. 2014; Matías-Hernández et al. 2016).

## Author contributions

N. Mahna and NFA conceptualized the research. AD, N. Maghsoudi, FAL, LMH and N. Mahna conducted the experiments. AD and NFA analyzed the data. AD and N. Mahna wrote the manuscript. N. Mahna, AM, SP and KF provided funds and lab facilities and supervised the research. All authors reviewed, edited, and approved the manuscript.

## 7. Supplementary Tables

**Supplementary Table 1.**
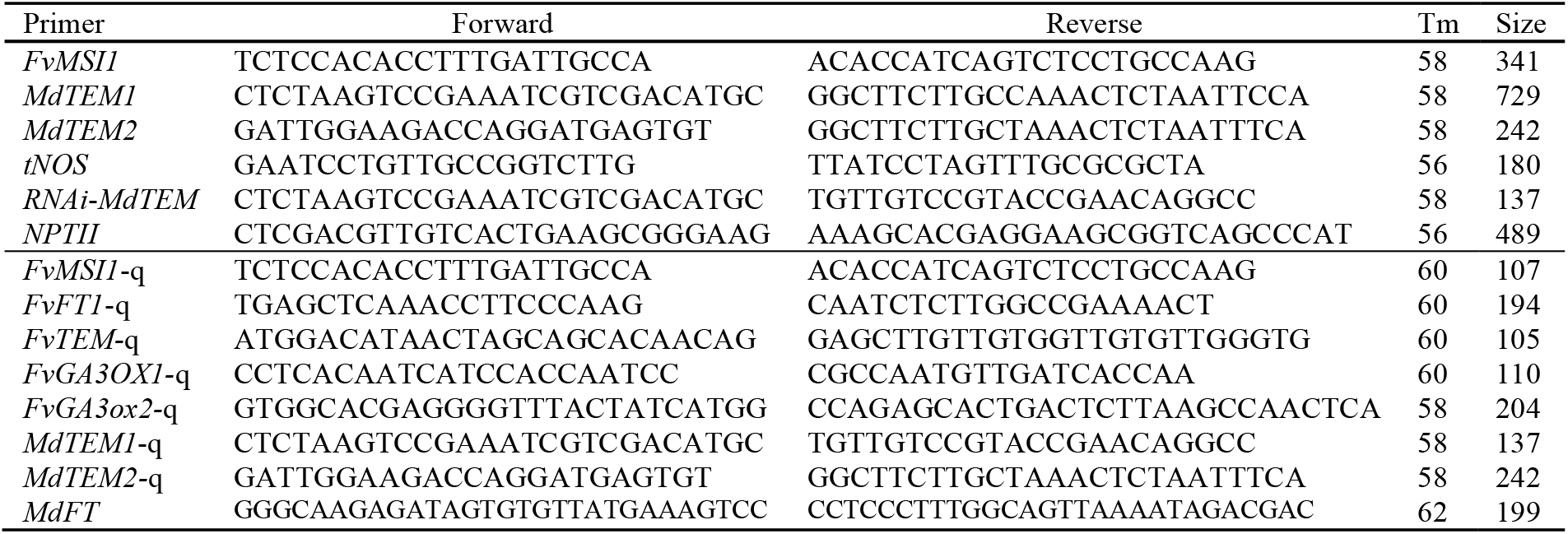
Primers used for PCR and quantitative Real-time PCR analyses

**Supplementary Table 2.** Transformation efficiency of wild strawberry based on the percentage of PCR-positive plants out of the total number of inoculated explants

| Construct | No. of Inoculated explants | No. of PCR positive plants | Transformation efficiency |
| --- | --- | --- | --- |
| 35S::*MdTEM1* | 140 | 33 | 23.57 |
| 35S::*MdTEM2* | 164 | 28 | 17.07 |
| RNAi-*tem* | 120 | 39 | 32.5 |

**Supplementary Table 3.** ANOVA of morphological flowering traits in wild strawberry

| S.O.V | df | Mean Square |  |  |
| --- | --- | --- | --- | --- |
|  |  | No. of leaves before flowering | No. of days before flowering | Trichome per mm<sup>2</sup> |
| Treatment | 3 | 86.48** | 300.54** | 447.37** |
| Error | 25 | 4.02 | 8.96 | 10.24 |
| C.V |  | 17.32 | 9.95 | 11.18 |

<sup>ns</sup> no significant, \*and \*\* significant at 5% and 1% probability level, respectively.

## 8. Supplementary Figs

**Supplementary Fig. 1.**
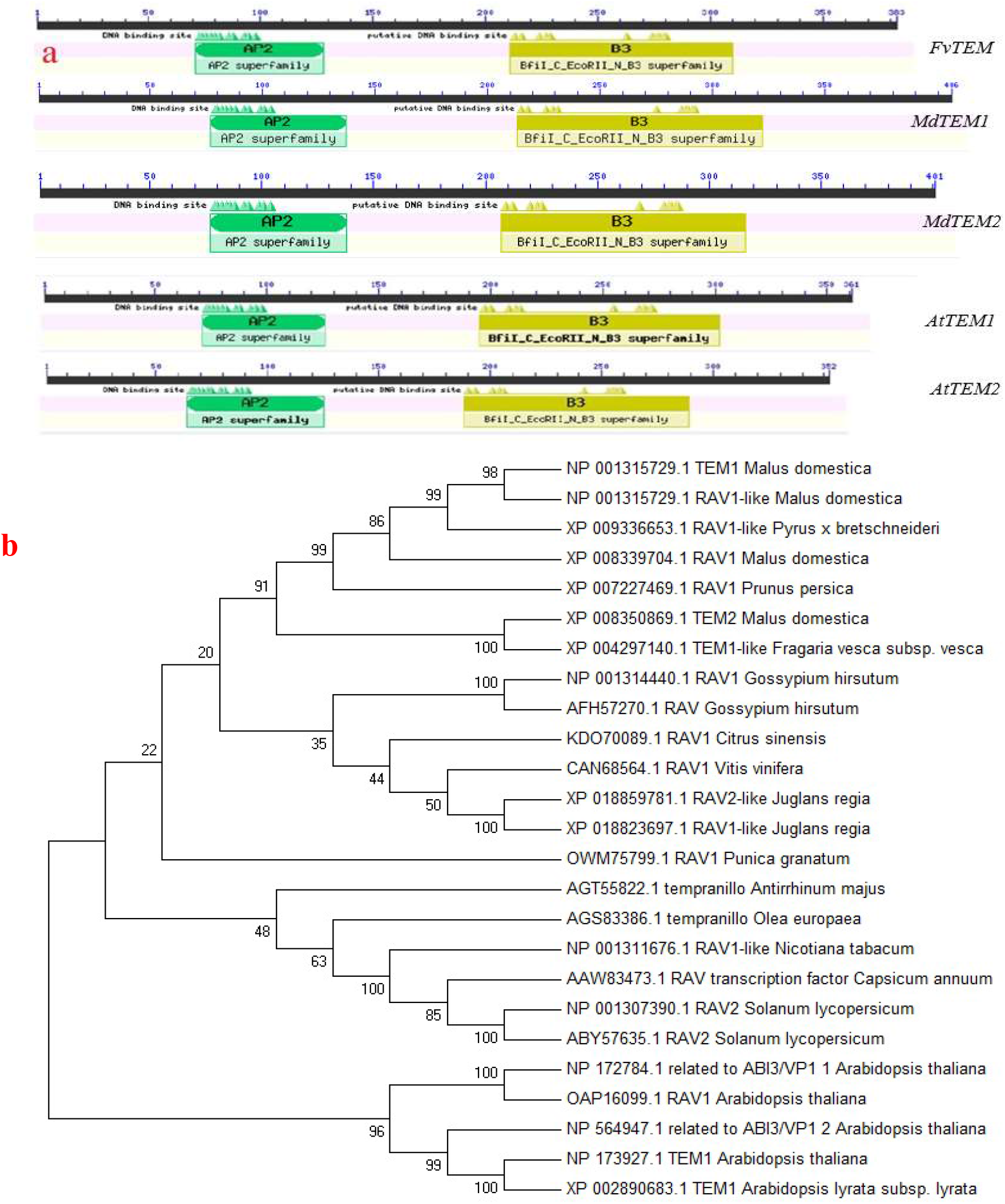
Phylogenetic analysis between MdTEM1/2, FvTEM and RAV sub-family class I members. a) Comparison of protein domain structure in FvTEM, MdTEM1 and MdTEM2, AtTEM1 and AtTEM2. b) Phylogenetic analysis of *FvTEM* deduced amino acid sequences and other RAV sub-family class I members homologs. The evolutionary history was inferred using the *Neighbor-Joining* method. The bootstrap consensus tree inferred from 500 replicates is taken to represent the evolutionary history of the taxa analyzed. The percentage of replicate trees in which the associated taxa clustered together in the bootstrap test (500 replicates) are shown next to the branches. Evolutionary analyses were conducted in *MEGA7*. Accession numbers are given next to the species name.

**Supplementary Fig. 2.**
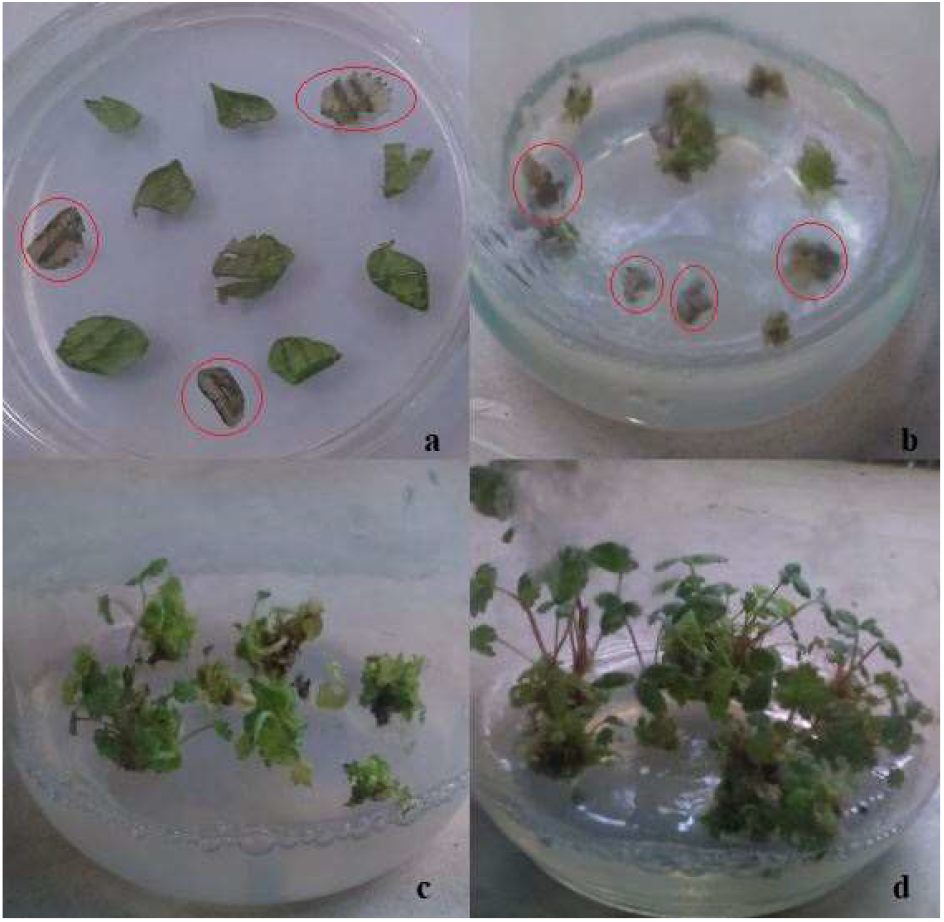
Selection and regeneration of putative transgenic lines of wild strawberry. a and b) Untransformed explants became necrotic (red), c and d) regeneration and proliferation of putative transformed explants.

**Supplementary Fig. 3.**
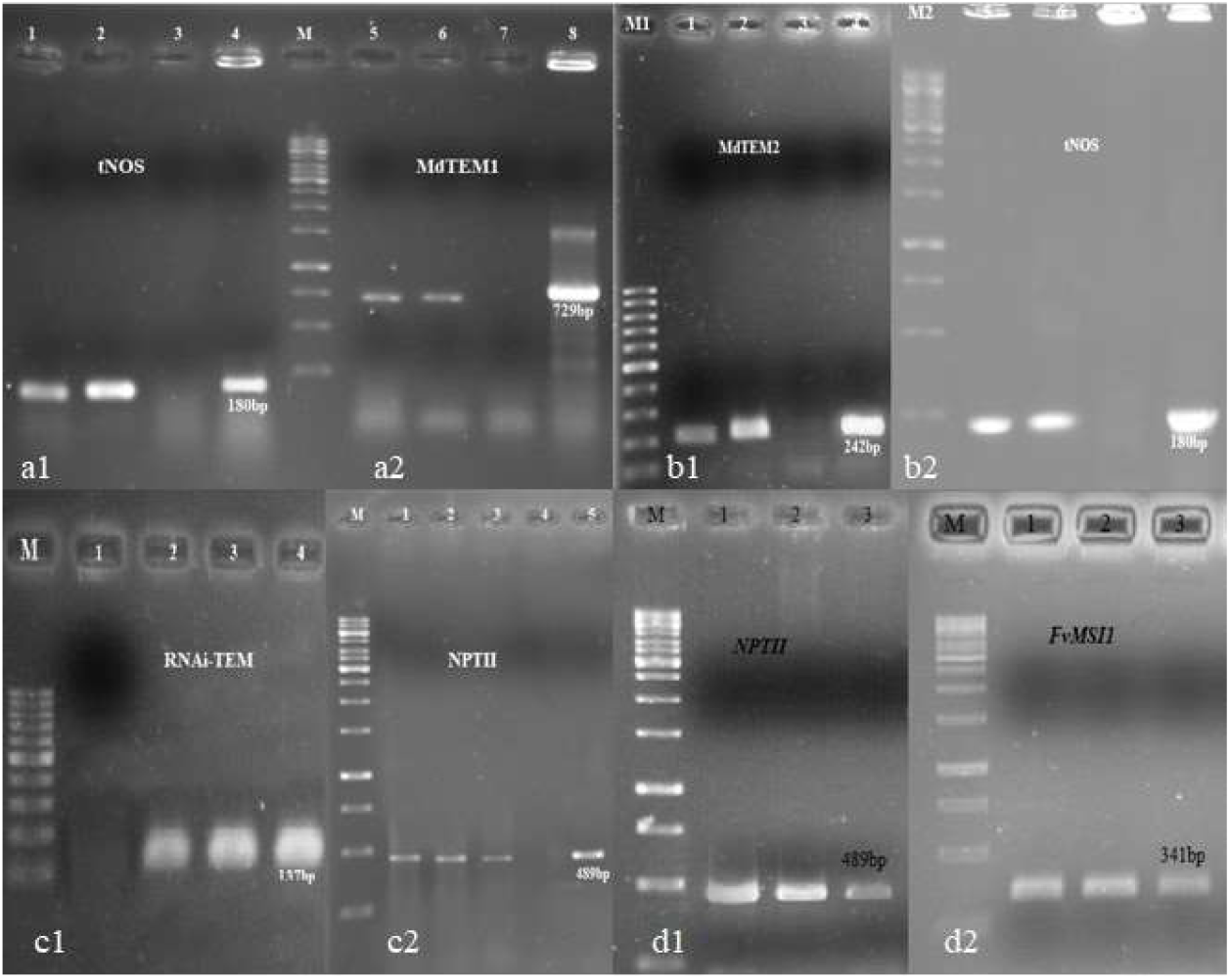
Wild strawberry plant transformation confirmation by PCR using tNOS, MdTEM1 and MdTEM2 primers for overexpression lines, and RNAi-tem and NPTII primers for RNAi silencing lines. a1 and a2: *35S::MdTEM1* (1 and 2: *35S::MdTEM1* lines, 3: control plant, 4: plasmid control); b1 and b2: *35S::MdTEM2* (1 and 2: *35S::MdTEM*2 lines, 3: control plant, 4: plasmid control); c1 and c2: RNAi-*MdTEM* (2,3 and 4 (c1) and 1,2,3 (c2): three RNAi-*MdTEM* lines); d1 and d2: RT-PCR of RNAi-*MdTEM* using NPTII and FvMSI1 primers.

## References

Aguilar-Jaramillo AE, Marín-González E, Matías-Hernández L, Osnato M, Pelaz S, Suárez-López P (2019) TEMPRANILLO is a direct repressor of the micro RNA miR172. The Plant Journal 100: 522–535

Bensmihen S, To A, Lambert G, Kroj T, Giraudat J, Parcy F (2004) Analysis of an activated ABI5 allele using a new selection method for transgenic Arabidopsis seeds. FEBS letters 561: 127–131

Brown T, Wareing P (1965) The genetical control of the everbearing habit and three other characters in varieties of Fragaria vesca. Euphytica 14: 97–112

Castillejo C, Pelaz S (2008) The balance between CONSTANS and TEMPRANILLO activities determines FT expression to trigger flowering. Current Biology 18: 1338–1343

Clough SJ, Bent AF (1998) Floral dip: a simplified method for Agrobacterium-mediated transformation of Arabidopsis thaliana. The plant journal 16: 735–743

Corbesier L, Vincent C, Jang S, Fornara F, Fan Q, Searle I, Giakountis A, Farrona S, Gissot L, Turnbull C (2007) FT protein movement contributes to long-distance signaling in floral induction of Arabidopsis. Science 316: 1030–1033

Dejahang A, Mahna N, Akhtar NF, Mousavi A (2018) Identification, isolation and in silico characterization of Fragaria vesca homologue of TEMPRANILLO gene. BIONATURE 38: 337–346

Doyle JJ, Doyle JL (1990) Isolation ofplant DNA from fresh tissue. Focus 12: 39–40

Esumi T, Tao R, Yonemori K (2005) Isolation of LEAFY and TERMINAL FLOWER 1 homologues from six fruit tree species in the subfamily Maloideae of the Rosaceae. Sexual Plant Reproduction 17: 277–287

Figueiredo AST, Resende JTV, Morales RGF, Gonçalves APS, Da Silva PR (2013) The role of glandular and non-glandular trichomes in the negative interactions between strawberry cultivars and spider mite. Arthropod-Plant Interactions 7: 53–58

Flachowsky H, Hanke MV, Peil A, Strauss S, Fladung M (2009) A review on transgenic approaches to accelerate breeding of woody plants. Plant Breeding 128: 217–226

Flachowsky H, Peil A, Sopanen T, Elo A, Hanke V (2007) Overexpression of BpMADS4 from silver birch (Betula pendula Roth.) induces early-flowering in apple (Malus× domestica Borkh.). Plant Breeding 126: 137–145

Folta KM, Davis TM (2006) Strawberry genes and genomics. Critical Reviews in Plant Sciences 25: 399–415

Folta KM, Gardiner SE (2009) Genetics and genomics of Rosaceae. Springer,

Fornara F, de Montaigu A, Coupland G (2010) SnapShot: control of flowering in Arabidopsis. Cell 141: 550–550.e552

Heide OM, Sønsteby A (2007) Interactions of temperature and photoperiod in the control of flowering of latitudinal and altitudinal populations of wild strawberry (Fragaria vesca). Physiologia Plantarum 130: 280–289

Huijser P, Schmid M (2011) The control of developmental phase transitions in plants. Development 138: 4117–4129

Jiao Y (2016) Trichome Formation: Gibberellins on the Move. Plant physiology 170: 1174–1175

Jung C (2017) Flowering time regulation: Agrochemical control of flowering. Nature plants 3: 17045

Koskela EA, Kurokura T, Toivainen T, Sønsteby A, Heide OM, Sargent DJ, Isobe S, Jaakola L, Hilmarsson H, Elomaa P (2017) Altered regulation of TERMINAL FLOWER 1 causes the unique vernalisation response in an arctic woodland strawberry accession. New Phytologist 216: 841–853

Koskela EA, Mouhu K, Albani MC, Kurokura T, Rantanen M, Sargent DJ, Battey NH, Coupland G, Elomaa P, Hytönen T (2012) Mutation in TERMINAL FLOWER1 reverses the photoperiodic requirement for flowering in the wild strawberry Fragaria vesca. Plant Physiology 159: 1043–1054

Kotoda N, Wada M (2005) MdTFL1, a TFL1-like gene of apple, retards the transition from the vegetative to reproductive phase in transgenic Arabidopsis. Plant Science 168: 95–104

Kotoda N, Wada M, Kusaba S, Kano-Murakami Y, Masuda T, Soejima J (2002) Overexpression of MdMADS5, an APETALA1-like gene of apple, causes early flowering in transgenic Arabidopsis. Plant Science 162: 679–687

Li C, Yamagishi N, Kasajima I, Yoshikawa N (2019) Virus-induced gene silencing and virus-induced flowering in strawberry (Fragaria× ananassa) using apple latent spherical virus vectors. Horticulture research 6: 18

Liu Y, Zhao Q, Meng N, Song H, Li C, Hu G, Wu J, Lin S, Zhang Z (2017) Over-expression of EjLFY-1 Leads to an Early Flowering Habit in Strawberry (Fragaria× ananassa) and Its Asexual Progeny. Frontiers in plant science 8: 496

Maghsoudi N, Mahna N, Sokhandan-Bashir N, Baghban-Kohnehrooz B (2015) Identification and isolation of a homolog of AtSEF gene from apple (Malus domestica). International Journal of Biosciences (IJB) 6: 153–161

Marín-González E, Matías-Hernández L, Aguilar-Jaramillo AE, Lee JH, Ahn JH, Suárez-López P, Pelaz S (2015) SHORT VEGETATIVE PHASE up-regulates TEMPRANILLO2 floral repressor at low ambient temperatures. Plant physiology 169: 1214–1224

Matías-Hernández L, Aguilar-Jaramillo AE, Marín-González E, Suárez-López P, Pelaz S (2014) RAV genes: regulation of floral induction and beyond. Annals of botany: mcu069

Matías-Hernández L, Aguilar-Jaramillo AE, Osnato M, Weinstain R, Shani E, Su P, Pelaz S (2016) TEMPRANILLO reveals the mesophyll as crucial for epidermal trichome formation. Plant physiology: pp. 01309.02015

Monte D, Somerville S (2002) Pine tree method for isolation of plant RNA. In: Bowtell D, Sambrook J eds. DNA microarrays: a molecular cloning manual. Cold Spring Harbor Laboratory Press, New York. pp. 124–126

Oosumi T, Gruszewski HA, Blischak LA, Baxter AJ, Wadl PA, Shuman JL, Veilleux RE, Shulaev V (2006) High-efficiency transformation of the diploid strawberry (Fragaria vesca) for functional genomics. Planta 223: 1219–1230

Osnato M, Castillejo C, Matías-Hernández L, Pelaz S (2012) TEMPRANILLO genes link photoperiod and gibberellin pathways to control flowering in Arabidopsis. Nature communications 3: 808

Pantazis CJ, Fisk S, Mills K, Flinn BS, Shulaev V, Veilleux RE, Dan Y (2013) Development of an efficient transformation method by Agrobacterium tumefaciens and high throughput spray assay to identify transgenic plants for woodland strawberry (Fragaria vesca) using NPTII selection. Plant Cell Reports 32: 329–337

Rantanen M, Kurokura T, Mouhu K, Pinho P, Tetri E, Halonen L, Palonen P, Elomaa P, Hytönen T (2014) Light quality regulates flowering in FvFT1/FvTFL1 dependent manner in the woodland strawberry Fragaria vesca. Frontiers in Plant Science 5:

Schmittgen TD, Livak KJ (2008) Analyzing real-time PCR data by the comparative C T method. Nature protocols 3: 1101

Sgamma T, Jackson A, Muleo R, Thomas B, Massiah A (2014) TEMPRANILLO is a regulator of juvenility in plants. Scientific Reports 4: 3704

Slovin JP, Schmitt K, Folta KM (2009) An inbred line of the diploid strawberry Fragaria vesca f. semperflorens for genomic and molecular genetic studies in the Rosaceae. Plant Methods 5: 1

Wada M, Cao Q-f, Kotoda N, Soejima J-i, Masuda T (2002) Apple has two orthologues of FLORICAULA/LEAFY involved in flowering. Plant Molecular Biology 49: 567–577

Wang L, Zhou CM, Mai YX, Li LZ, Gao J, Shang GD, Lian H, Han L, Zhang TQ, Tang HB (2019) A spatiotemporally regulated transcriptional complex underlies heteroblastic development of leaf hairs in Arabidopsis thaliana. The EMBO journal 38: e100063

Wang S, Normal N, Zeng Q, Wang Y, Pattanaik S, Yuan L (2014) An overview of the gene regulatory network controlling trichome development in the model plant, Arabidopsis. Regulation of Cell Fate Determination in Plants: 12

Wellmer F, Riechmann JL (2010) Gene networks controlling the initiation of flower development. Trends in Genetics 26: 519–527

Yamagishi N, Kishigami R, Yoshikawa N (2014) Reduced generation time of apple seedlings to within a year by means of a plant virus vector: a new plant-breeding technique with no transmission of genetic modification to the next generation. Plant biotechnology journal 12: 60–68

